# Concurrent Contribution of Co-contraction to Error Reduction during Dynamic Adaptation of the Wrist

**DOI:** 10.1101/2022.10.26.513872

**Authors:** Andria J. Farrens, Kristin Schmidt, Hannah Cohen, Fabrizio Sergi

## Abstract

MRI-compatible robots provide a means of studying brain function involved in complex sensorimotor learning processes, such as adaptation. To properly interpret the neural correlates of behavior measured using MRI-compatible robots, it is critical to validate the measurements of motor performance obtained via such devices. Previously, we characterized adaptation of the wrist in response to a force field applied via an MRI-compatible robot, the MR-SoftWrist. Compared to arm reaching tasks, we observed lower end magnitude of adaptation, and reductions in trajectory errors beyond those explained by adaptation. Thus, we formed two hypotheses: that the observed differences were due to measurement errors of the MR-SoftWrist; or that impedance control plays a significant role in control of wrist movements during dynamic perturbations. To test both hypotheses, we performed a two-session counterbalanced study. In both sessions, participants performed wrist pointing in three force field conditions (zero force, constant, random). Participants used either the MR-SoftWrist or the UDiffWrist, a non-MRI-compatible wrist robot, for task execution in session one, and the other device in session two. To measure anticipatory co-contraction associated with impedance control, we collected surface EMG of four forearm muscles. We found no significant effect of device on behavior, validating the measurements of adaptation obtained with the MR-SoftWrist. EMG measures of co-contraction explained a significant portion of the variance in excess error reduction not attributable to adaptation. These results support the hypothesis that for the wrist, impedance control significantly contributes to reductions in trajectory errors in excess of those explained by adaptation.

## I. Introduction

Neurorehabilitation is centered on the idea that retraining motor function can be advanced by incorporating concepts of neuromotor control into therapy. Although many every day manipulation tasks are performed using the hand and wrist, relatively few studies have focused on the neuromotor control of the wrist, especially during human-robot interaction. To address this gap in knowledge, our group has developed the MR-SoftWrist, an fMRI-compatible 3 degree of freedom (DOF) wrist robot, to study neuromotor control of the wrist during sensorimotor tasks via fMRI [1], [2]. In our work, we focus on characterizing adaptation-specific processes, which enable our flexible control of movement in novel dynamic environments, and contribute to the motor learning process.

Adaptation refers to the formation of an internal model of the task dynamics, learned through trial and error, that is used to predict and compensate for environmental perturbations [3], [4]. Adaptation results in transient changes in behavior (after effects) when perturbations are removed. To localize brain regions responsible for this dynamic motor control process, we seek to associate measurements of motor kinematics and kinetics with measurements of brain function obtained via fMRI [5]. Previously, adaptation data collected via the MR-SoftWrist showed some differences with those measured during arm reaching tasks in the adaptation literature [6], [7]. First, the magnitude of adaptation was lower than what is typically reported for reaching tasks following a similar number of trials [8], [9]. Moreover, the end magnitude of adaptation observed in our studies did not explain the large reduction in trajectory errors observed.

From these results, we formed two hypotheses. The first was that behavioral measures taken with the MR-SoftWrist may be influenced by the inherent compliance of the device. Adaptation is measured during error clamp trials, in which the robot applies a force tunnel that maximally restricts lateral deviations to measure participants force profiles that reflect their prediction of required task dynamics. Ideally, the walls of this force tunnel would be infinitely stiff to enable perfect measurement of the applied forces. To achieve MRI-compatibility, the MR-SoftWrist utilizes a series elastic actuator design with piezoelectric motors that results in a limited error clamp stiffness bounded by the stiffness of the springs [1], [10]. This inherent compliance of the MR-SoftWrist makes it unclear if the differences in adaptation observed in our experiments are due to differences in motor control of the wrist, or due to limitations in the measurement of force profiles taken with the MR-SoftWrist. To address this issue, we developed a benchtop wrist robot, the UDiffWrist, that can apply virtual walls with much greater stiffness to validate our behavioral measures of dynamic adaptation of the wrist executed with the MR-SoftWrist [11].

Alternatively, if our behavioral measures are not unduly influenced by measurement error, we hypothesized that reduction in trajectory errors in excess of those predicted by our measures of adaptation were due to concurrent co-contraction. All behavioral measures analyzed in our study of adaptation focus on the feed-forward phase of movement. In the feed-forward phase of movement, impedance control is known to contribute to error reduction in novel or unlearnable dynamic environments [12]–[15]. Impedance control refers to the generalized strategy of stiffening the joint via muscle co-contraction to reduce kinematic errors caused by perturbations, and can be assessed via EMG. In arm reaching tasks, impedance control is typically dominant early in task execution, but is reduced to negligible amounts as adaptation progresses. From our previous studies, it appears that impedance control may play a more persistent role in control of the wrist, contributing to excess error reduction not explained by measures of adaptation alone.

In this study, we tested both hypotheses via a two-session counterbalanced experimental design. In both sessions, participants performed the same task schedule, consisting of a zero force task, an adaptation task, and a random force task. Half the participants used the MR-SoftWrist for task execution in session 1, while the other half used the UDiffWrist. For session 2, participants used the other robotic device for task execution. To limit effects of savings on session 2 behavior, sessions 1 and 2 were conducted 3-4 weeks apart. For all tasks, muscle activation of the primary wrist muscles engaged in task execution—flexor carpis radialis (FCR), flexor carpis ulnaris (FCU), extensor carpis radialis (ECR), extensor carpis ulnaris (FCU)— was collected using surface EMG [16].

To validate behavioral measures taken with the MR-SoftWrist, we compared behavior measured between devices and experimental sessions to test the hypothesis that there would be no significant differences between robotic devices. To investigate concurrent co-contraction, we compared EMG measured in the dynamic adaptation task, in which participants are expected to use a predominantly adaptation-based strategy for error reduction, to EMG measured in the random force task, in which we expect participants to use a predominantly co-contraction-based strategy for perturbation rejection. To investigate the contributions of co-contraction to excess error reductions, we hypothesized that error reduction in excess of that explained by models of neuromotor adaptation would be associated with EMG measures of co-contraction.

## II. Materials and Methods

Our study utilized a two-session counter balanced design. In session 1, participants interacted with either the MR-SoftWrist or the UDiffWrist to perform a series of three wrist pointing tasks; a zero force task for characterization of baseline motor performance, a velocity-dependent curl force task for characterization of dynamic adaptation, and a randomly alternating curl force task for characterization of impedance control (Fig. 1). Approximately three weeks after session 1, participants performed the same set of tasks using the other robotic device in session 2. In each session, participants kinetics and kinematics were recorded by the robotic device used for task performance, while surface EMG of the four primary wrist muscles was acquired to measure muscle activity associated with neuromotor control in each task.

**Fig. 1.**
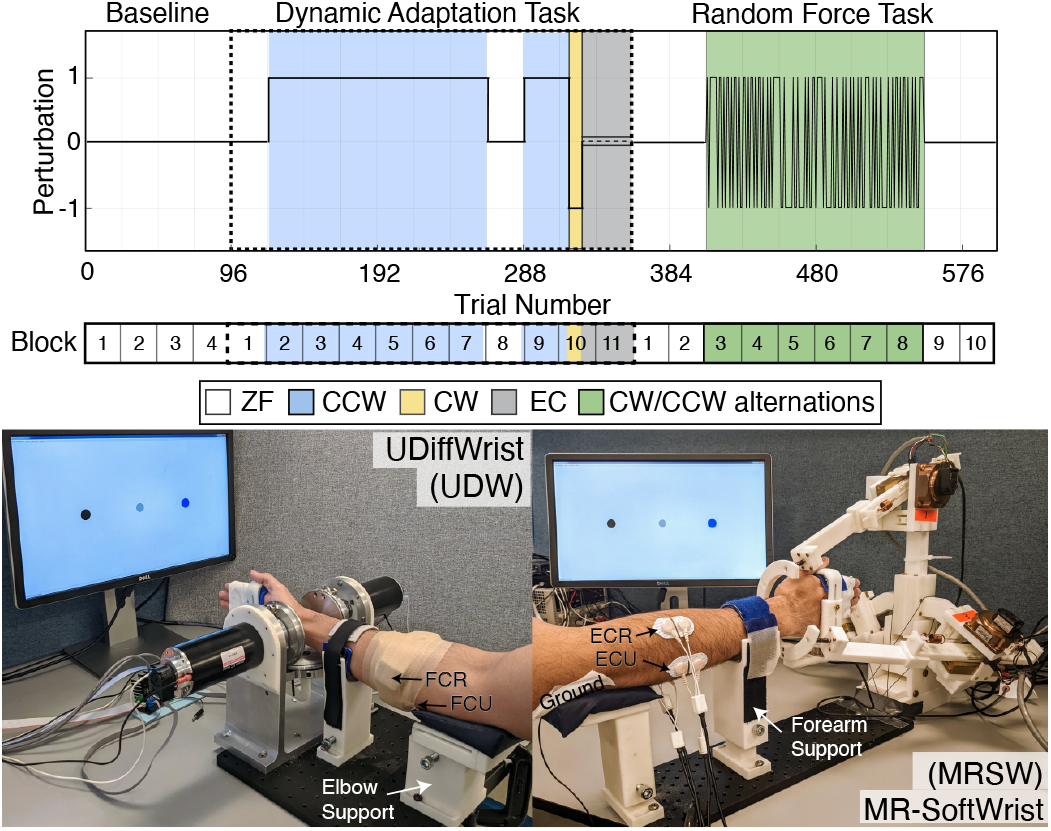
Experimental protocol. Top: Schematic depicting the three motor tasks performed in each session. The three tasks are divided by dashed lines, and include the “baseline” motor performance task (4 blocks of 24 trials), the dynamic adaptation task (11 blocks of 24 trials), and the random force task (10 blocks of 24 trials). The solid black line depicts the magnitude and direction of the force field applied in each task (0 for zero force, 1 for CCW, −1 for CW). Blocks within each task are highlighted below the task schematic, for reference in definitions of experimental phases of interest used in our statistical analyses. Bottom: Experimental set up displaying the robotic devices, handle and forearm supports, and associated EMG systems. The UDiffWrist is shown with the wireless Delsys EMG system, and the MR-SoftWrist is shown with the cabled OT-Bioelettronica system.

### A. Participants

27 healthy young adults free from neurological or musculoskeletal injury participated in this study. 15 participants were assigned to use the MR-SoftWrist for session 1 and the UDiffWrist for session 2 (MRSW-UDW), and twelve participants were assigned the reverse order (UDW-MRSW). One participant from each group was eliminated for failure to follow task instructions, and one participant from the MRSW-UDW group was removed for hardware malfunction, resulting in 13 individuals with session 1 data for the MR-SoftWrist (N = 13; 6 male, age: 25 ±3.58 years), and 11 individuals for the UDiffWrist (N = 11; 5 male, age: 25 ±3.67 years). 4 participants from MRSW-UDW group, and 2 participants from the UDW-MRSW group did not return for the second session, resulting in 9 participants in each group with data for both sessions. The study was approved by the Institutional Review Board of the University of Delaware, IRB no 906215-10.

### B. Experimental Procedure

The experimental protocol is shown in Fig. 1. Prior to task performance, EMG sensors were placed on the participant, and appropriately sized handle and forearm supports were selected (Fig. 1, Bottom). Participants used the same handle and support for both robotic devices. Participants’ hands were secured into the handle such that it fit snug around their open palm, without touching the fingers. Participants’ fingers were taped with athletic tape at the finger joints to prevent grasping of the handle. For each session, participants performed a familiarization task with the robotic device to learn task instructions and interaction with the robot in the zero force condition. Following training, participants performed a series of three motor tasks over roughly 40 minutes. EMG activation was measured during isometric contractions performed immediately before the familiarization task and after the last task for normalization purposes and to assess any changes in signal quality across the session. Participants wore noise cancelling headphones (COWIN E7) that played white noise to eliminate environmental distractions during task execution.

### C. Robotic Devices

The MR-SoftWrist is an fMRI-compatible robot (Fig. 1, Bottom right) that supports flexion-extension (FE) and radial-ulnar deviation (RUD) on the wrist in a circular workspace (radius 20 deg). The device has a maximum output torque of 1.5 N·m about each axis, and can display kinesthetic environments ranging from a zero force mode that minimally perturbs the user’s movements (trial average resistive torque = 0.05 Nm; peak resistive torque < 0.17 Nm), to a stiffness control mode that displays a high stiffness environment (max virtual stiffness, *k_v_* = 0.45 Nm/deg) [1]. User interaction forces are measured via springs and linear encoders placed in series between the piezoelectric motors and the load (user).

The UDiffWrist (Fig. 1, Bottom left) is a low-impedance wrist robot that features a cable-differential transmission with a workspace matching the one of the MR-SoftWrist. The device has a maximum output torque of 31.4 Nm about each axis (FE, RUD), and a lower impedance compared with the MR-SoftWrist [11]. For all tasks, the impedance of the UDiffWrist was increased via application of a viscous damping force to match that of the MR-SoftWrist, confirmed via experiments performed in a zero force condition. User interaction forces were measured using a 6-axis force transducer ATI Mini40 F/T Sensor located at the base of the handle.

In this experimental protocol, each robot operated in one of three control modes: a zero force mode (ZF), a curl force mode (CF), and an error clamp mode (EC). In the zero force mode, the desired interaction torque is set to zero (*τ_FE_* = *τ_RUD_* = 0) to display a transparent environment to the user. In the curl force mode the robot applied a velocity-dependent torque defined as 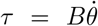 such that *τ* = [*τ_FE_, τ_RUD_*], proportional to the measured velocity 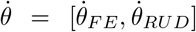. Clockwise and counter-clockwise force fields were achieved with B = ± 3.49 Nm·s/deg, respectively [3]. Finally, in the error clamp mode, the robot produced a force channel that clamps lateral trajectory errors to zero, to measure participants lateral force profiles that reflect their expectation of required task dynamics [10].

The MR-SoftWrist implemented error clamp forces with a positional deadband of 0.025 deg, a stiffness of 500 mNm/deg and damping of 5 mNm·s/deg, and a saturation of 1 Nm. The UDiffWrist implemented error clamp trials with a deadband of 0.0025 deg, a stiffness of 1250 mNm/deg and damping of 15 mNm·s/deg, with a saturation of 1.2 Nm. Values for error clamp force fields were tuned via trial and error to achieve the stiffest possible virtual walls given the capabilities of the hardware, as typical in these studies. For the MR-SoftWrist, error clamp trials restricted maximal lateral deviations to (mean ±std) 0.205 ±0.037 deg across all dynamic conditions, with a range of [−0.394, 0.309] deg across participants. For the UDiffWrist, error clamp trials restricted lateral errors to 0.0847 ±0.021 deg across dynamic conditions, with a range of [−0.312, 0.136] deg across participants. Due to inherent hardware limitations of the MR-SoftWrist, the UDiffWrist achieved a greater restriction of lateral deviation, although both devices sufficiently clamped errors close to zero.

For both devices, gravity compensation and force control were implemented at 1000 Hz via Simulink controllers using the Quarc real-time engine coded in MATLAB 2017a. Interaction force, velocity and position data were logged at a rate of 1000 Hz using a on Quanser Q8-USB data acquisition card. For more on the control and design of the MR-SoftWrist, see [1], and for the UDiffWrist see [11].

### D. EMG

At the beginning of each session, the four muscles of the wrist—flexor carpis radialis (FCR), flexor carpis ulnaris (FCU), extensor carpis radialis (ECR), extensor carpis ulnaris (FCU)—were identified via palpation of the forearm during muscle contraction (wrist flexion-extension and radial-ulnar deviation). To find the optimal position for sensor placement, the electrode was moved along the muscle belly until the largest burst of activation was measured during muscle contraction. The location then prepared for electrode placement by shaving the arm hair, and cleaning the skin with abrasive electrode gel followed alcohol.

Due to differences in electromagnetic interference generated by the motors of each robot, we used two different EMG systems to acquire muscle activation. For the MR-SoftWrist, we used an OT Bioelettronica amplifier and software (sampling frequency: 2048 Hz), and for the UDiffWrist we used a Delsys wireless electrode system and software (sampling frequency: 2000 Hz). For the OT Bioelettronica system, the common ground electrode was placed at the elbow. For both systems, EMG signal was time synced to task performance data via a common analog signal. Electrodes were not moved or replaced between tasks.

At the beginning and end of each session, EMG activation was measured during isometric contractions performed in the robotic device used for task execution. With the end effector of the robot locked in place, participants were cued to apply and hold a torque of 1 Nm for 7 seconds twice in each direction (flexion, extension, radial deviation, and ulnar deviation) to confirm electrode placement and signal quality. Forces generated by the participant were measured by a force torque sensor (ATI Mini40 F/T Sensor) placed underneath the handle. A force level of 1 Nm was chosen as it elicited clean bursts of activation, while remaining on roughly the same scale as the peak force expected in our task (anticipated ~ 400 mNm for expected velocities within our cued trial duration).

### E. Paradigm

Each session included three motor tasks: a motor performance task, a dynamic adaptation task, and a random force task (Fig. 1, Top). In each task, participants moved their wrist to control a cursor displayed on a monitor (Fig. 1, Bottom). The cursor was displayed continuously as a grey circle (radius 1 deg). Flexion-extension of the wrist moved the cursor horizontally, while radial-ulnar deviation moved the cursor vertically. Pronation-supination was prevented by a forearm support. Participants were cued to make alternating flexion-extension rotations to move a cursor in a straight line to one of two circular targets (radii 1.25 deg) located at (±10, 0) degrees in flexion-extension, radial-ulnar deviation.

Trial onset was cued by a change in target color from black to blue. Trial completion was achieved when the error between the cursor and the target was less than 1.5 deg for greater than 250 ms. After trial completion, the reached target provided timing feedback for 0.5 s by turning red if the movement duration was greater than 650 ms or green if it was less than 300 ms. Otherwise, the target remained black. The inter-trial-interval between timing feedback and cuing of the next trial was randomly selected from a normal distribution *N*(1.25, 0.2) s, bounded between [0.25 1.75] s. Each task was divided into blocks of 24 trials, with 8 s rest periods between blocks to prevent fatigue.

The motor performance task consisted of 4 blocks (96 trials) performed in the zero force mode to measure behavior (kinetics, kinematics, EMG) during unperturbed wrist pointing.

The dynamic adaptation task consisted of 11 total blocks. The first 8 blocks of the task consisted of an initial block of zero force trials, followed by 6 blocks in a counter-clockwise curl force field, followed by a block of zero force trials. This portion of the task was used to assess adaptation to a constant perturbation and immediate after effects of training, consistent with our previous studies. Blocks 9-11 followed an A-B-EC perturbation schedule, that consisted of 30 counter-clockwise trials, 10 clockwise trials, and 32 error clamp trials. This A-B-EC perturbation schedule, in which participants re-adapt to an initial perturbation (A, counter-clockwise), followed by brief exposure to an oppositely directed perturbation (B, clockwise), before having their errors clamped to zero (EC), should elicit a rebound of behavior (spontaneous recovery) that reflects adaptation to the first perturbation (A), and produce unique model parameter estimates [6], [9].

The random force task consisted of 10 blocks; two initial blocks of zero force trials to reestablish “baseline” behavior, 6 blocks in an alternating curl force condition, and two final blocks in a zero force condition to enable assessment of any after effects. The alternating curl force field force was generated in a pseudo-random manner. Force-field direction changed with a 50% probability between each trial, and logic was applied to prevent greater than 4 consecutive trials occurring in a ‘constant’ field. For this purpose, we defined ‘constant’ fields as conditions in which all trials were clockwise, counter-clockwise, or in which alternations occurred on every trial sequentially (i.e. CW-CCW-CW-CCW) resulting in the application of a constant upwards or downward force. Clockwise and counter-clockwise perturbations were evenly balanced across this task, such that the average force experienced across the task was 0.

Error clamp trials were interspersed pseudo-randomly throughout each task with a 1/8th probability, except for the B (no error clamp trials) and EC (all error clamp trials) portion of the dynamic adaptation task [10]. Error clamp trials only occurred after the first two trials of a block, and had between trial spacing of 3 to 12 trials. Across devices and sessions, all participants performed the exact same task schedule.

### F. Data Analysis

#### 1) Data Preprocessing

Processing of behavioral data was conducted using MATLAB (The MathWorks, version 2020b). All position, velocity, and force data were low-pass filtered with a zero-shift 4th order Butterworth filter with a 25 Hz cut off frequency using MATLAB’s filtfilt function. For each trial, position and force data from trial onset to trial end were resampled into 1000 data points, and divided between extension (right) and flexion (left) movements. For both sessions, baseline trajectory and force profiles were determined for each direction as the average of all valid trials from blocks 2-4 of the motor performance task and block 1 of the dynamic adaptation task. Within each session, direction-specific baseline profiles were subtracted from the respective flexion- or extension-directed trials for force and position data, such that all behavioral metrics reflect a change relative to typical, non-perturbed wrist pointing.

For trial-by-trial data analysis, trial onset was defined as the instant the absolute cursor velocity exceeded 15 deg/s. Trial end was defined as the instant the cursor was within 3 deg of the target in the flexion-extension direction. Trials were excluded if they matched any of the following conditions: 1) a trial duration outside the [200, 700] ms range; 2) a maximum velocity below 40 deg/s; 3) a reversal in goal directed velocity that occurred before the trial max velocity, indicative of false starts. At the group level, this resulted in an average exclusion of 1 ±0.56% of all trials with a max removal of 2% (24 trials) across both sessions.

For EMG analysis, we used only session one data, which had the largest number of participants with usable data. EMG data were bandpass filtered at 30-500 Hz, rectified, and the envelope was taken via a low-pass zero-shift 4th order Butterworth filter with a 10 Hz cut off frequency [15]. EMG for each muscle was normalized by the average activation measured during isometric contractions in their preferred directions (4 repetitions measured per direction). This included activations measured in flexion and radial deviation for the FCR, in flexion and ulnar deviation for the FCU, in extension and ulnar deviation for the ECU, and extension and radial deviation for the ECR.

EMG data collected from four participants were excluded due to issues of signal drop out or significant drift across all muscle groups. For the ECU, the remaining 20 participants (MRSW, N = 12, UDW, N = 8) were included in all subsequent analyses. For the FCR, an additional participant was removed for low SNR, and two for signal drift, resulting in 17 total participants (MRSW, N=9, UDW, N= 8). For our global activation measure (detailed below), we included all participants (N = 20) with valid muscle activation measured in at least two of the four muscle groups.

#### 2) Behavioral outcome measures

All behavioral metrics were calculated in the first 150 ms after movement onset, to capture behavior associated with feed-forward motor-control processes that most directly reflect participants pre-planned motor actions [15], [17].

##### a) Adaptation index

As error clamp trials occur unexpectedly, force profiles measured in these trials reflect the participants’ expectation of the task dynamics [10]. Thus, we defined adaptation index on error clamp trials as the ratio between the area under the measured force profile and the area under the ideal force profile necessary to compensate for a counter-clockwise (CCW) curl force field [6]. An adaptation index of one indicates perfect adaptation to a CCW curl force field, while an adaptation index of negative one indicates perfect adaptation to a CW curl force field. An adaptation index of zero represents no adaptation.

##### b) Trajectory error

On all field trials, we calculated participants’ angular trajectory error (i.e. performance error) defined as the internal angle between the lines a) connecting the start target and the cursor at its maximum lateral deviation and b) connecting the start and end targets [18].

##### c) Excess error reduction

Based on the results of our previous work, we chose to use the two-state model of adaptation to explain behavior attributable to dynamic adaptation in our task [6], [8]. The model equations are reported below:

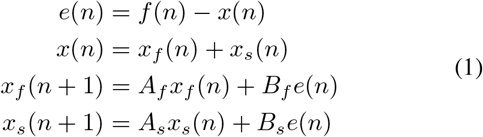

In this model, performance error, *e*(*n*), is modeled as the difference between the applied force field, *f*(*n*), and the participant’s motor output, *x*(*n*). *x*(*n*) represents adaptation of the internal model on trial *n*, that is quantified experimentally by adaptation index, and is the sum of two inner states; a fast learning state, *x_f_*, and a slow learning state, *x_s_* that are updated on every trial by performance error, *e*(*n*), in proportion to their retention (*A_f_* and *A_s_*) and update parameters (*B_f_* and *B_s_*). Here, *A_f_* < *A_s_* and *B_f_* >*B_s_*, as the slow state retains more from trial to trial and is less sensitive to error than the fast state. All parameters are constrained between (0, 1).

We hypothesized that reductions in trajectory errors *in excess* of those predicted by measures of adaptation were due to concurrent co-contraction. To define “excess error reduction”, we first fit the two-state model to adaptation index data measured in the dynamic adaptation task. Models fit to adaptation index data produce estimates of trajectory errors that are scaled by the magnitude of the perturbations experienced (1 for CCW/-1 for CW). To scale measured trajectory errors to model-estimated trajectory errors, we regressed the measured trajectory errors onto the model-estimated trajectory errors at each change in force application (i.e. trials 121, 265, 289, 319). We chose to use the initial trial following each removal/application of force as these transitions occur unexpectedly and most readily reflect behavior based on participants prediction of expected task requirements. Measured trajectory errors were then normalized by the scalar value returned by this regression.

For each participant, we quantified “excess error reduction” as the average residual error between their model-estimated trajectory errors (fit to subject-level adaptation index data) and normalized trajectory errors measured in the steady-state phase of the dynamic adaptation task (block 7), where we previously observed the largest discrepancy between adaptation and error reduction. At the group level, we defined excess error reduction as the squared residual error between model-estimated trajectory errors (fit to group average adaptation index data) and the normalized average trajectory errors measured across all tasks.

#### 3) EMG outcome measures

On each trial, we quantified the average EMG activation measured over the time interval of −150 ms to 50 ms surrounding movement onset. This time period was chosen based on previous studies that used intervals ranging from −200 ms to 130 ms surrounding movement onset to measure feed-forward muscle activation associated with co-contraction and adaptation, while avoiding influence from online error correction processes [15], [19], [20]. Visual inspection of the EMG traces showed that the time interval from −150 ms to 50 ms captured task related increases in muscle activation with limited influence from reflex related activations (Fig. 2, Top). Within this time window, the maximal average deviation was less than 1 degree, and occurred at 50ms (Fig. 2, bottom left). As short latency reflexes typically occur 30-50 ms following a muscle stretch, with long-latency reflexes occurring >50 ms post-stretch, the selected time window is expected to have limited influence from stretch related reflex activity. There was no significant difference in the absolute velocity measured within this time interval between task conditions (Fig. 2, bottom right).

**Fig. 2.**
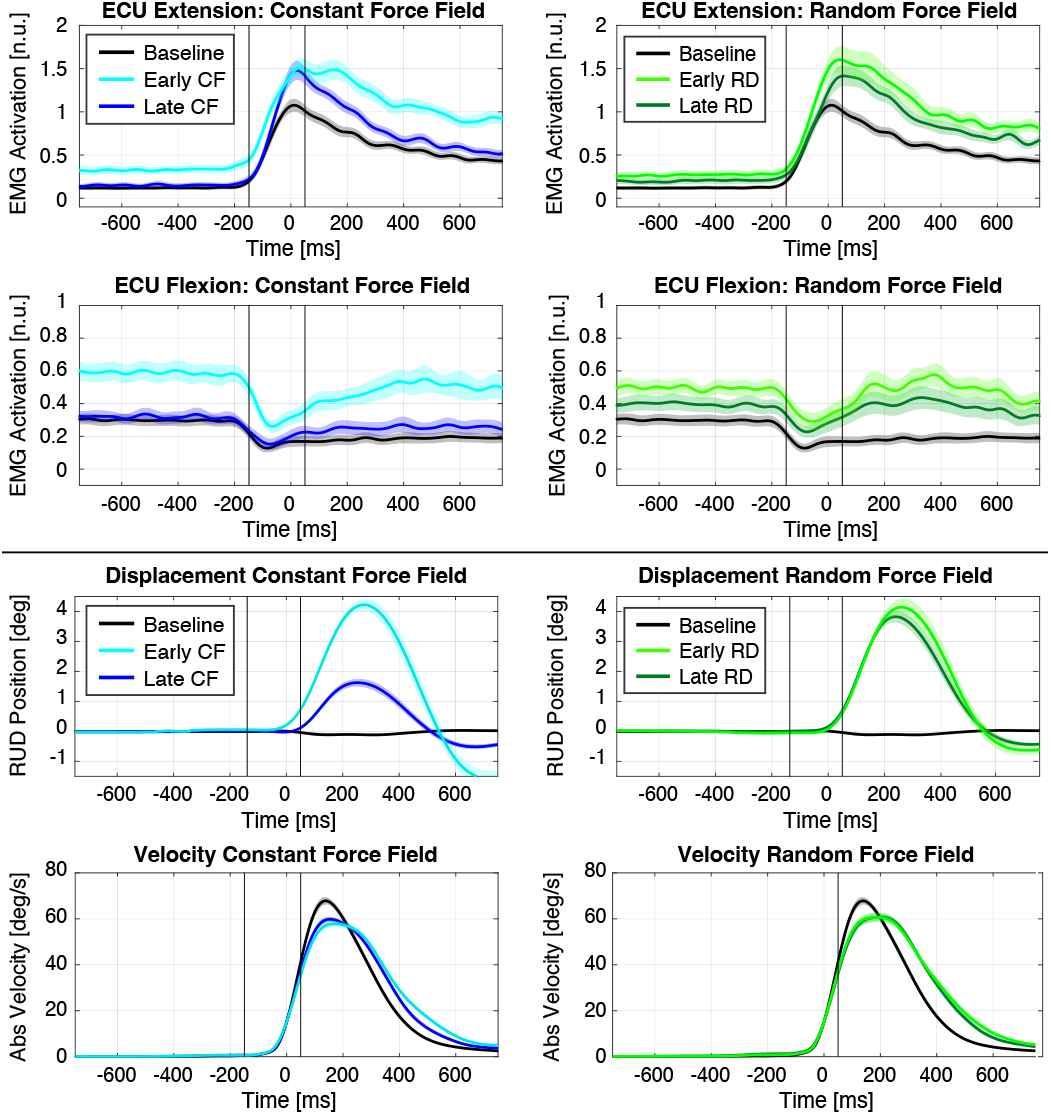
Average EMG and kinematic data measured in each task condition. Baseline included the average of all trials within the zero force condition. Early and Late force conditions (CF: Curl force, RD: Random force) depict the average of the first 24 and last 24 trials measured within each force condition, within the constant perturbation phase. EMG activation for the ECU muscle during extension (agonist) and flexion (antagonist) movements. Evaluation of lateral deviations and absolute velocity show task kinematics are well matched across conditions within the selected time window.

##### a) Antagonist muscle activation

To quantify co-contraction, we quantified activation measured in *pure antagonist* muscles to goal-directed movement. Previous studies have used “wasted contraction” to quantify antagonist muscle activation, quantified as the minimum activation measured between agonist and antagonist muscle pairs [20]. However, this measure requires the combination of signal from multiple muscles that may be influenced by differences in normalization and noise between EMG channels. As such, we chose to quantify activity of antagonist muscles using a univariate analysis simplified by the mechanics of our specific tasks.

In the zero force condition, both the ECR and ECU muscles act as antagonist muscles to goal-directed movement for flexion trials; similarly, for extension trials the FCR and FCU muscles act as antagonist muscles. In the dynamic adaptation task, in which a CCW force is applied, extension trials (right directed) experience radial (upwards) perturbations, while flexion trials (left directed) experience ulnar (downwards) perturbations. Consequently, for extension trials, we identified the FCR as a *pure* antagonist muscle, i.e. a muscle that acts as an antagonist both for goal-directed movement and perturbation rejection. Similarly, we identified the ECU as the pure antagonist muscle for flexion trials.

In the random force task, the perturbations alternated evenly between clockwise and counter-clockwise directions. Consequently, across the task, no muscle can be defined as a ‘pure’ agonist/antagonist in the radial-ulnar task dimension. However, given the balanced, rapid nature of these alternations, we do not expect this task to produce a learned response in any muscle group in either the ulnar or radial direction, but rather an increased activation along both dimensions evenly [21]. Moreover, the time window evaluated in our task occurs prior to significant reflex related activity associated with curl force perturbations that may influence muscle activation measured between tasks. As such, antagonist muscles in this task were defined based on goal-directed movement only, in the same way as the zero force condition. To remain consistent between tasks, we chose to examine activation in the FCR for extension trials and the ECU for flexion trials.

##### b) Global Activation

To enable comparisons with previous work, we additionally investigated global activation measured across all muscles as a proxy for stiffness [15], which accounts for activation of all agonist and antagonist muscles.

##### c) EMG measures of co-contraction

To investigate the association between EMG measures of co-contraction (antagonist, global) and excess error reduction, we quantified change in co-contraction for each participant as the difference between activation measured in the steady-state phase of the dynamic adaptation task (block 7) from their average baseline activation (blocks 3 and 4 of the motor performance task). At the group level, we quantified average co-contraction in antagonist muscles across all subjects for each trial for investigation of group level excess error reduction across all tasks.

#### 4) Statistical testing

##### a) Mixed model analysis of behavior

To test for differences in adaptation measured between devices we focused on the dynamic adaptation task, in which we expect participants to show maximal adaptation. To test for the effects of training on behavior, we used a mixed model with primary outcome measure of adaptation index. The between-subject factors were robotic device (levels: MR-SoftWrist, UDiffWrist), and session of task execution (levels: Session 1, Session 2). The within-subject factor was experimental phase, with levels: Baseline, defined as the last block of the motor performance task, Early Counter-Clockwise and Late Counter-Clockwise, defined as the beginning (block 2) and end (block 7) of the constant perturbation phase in the dynamic adaptation task, respectively, After Effects, defined as block 8 of the dynamic perturbation task performed in the zero force condition, Relearning, defined as the re-introduced counter-clockwise curl force condition spanning blocks 9 and 10, Initial Error Clamp, defined as the first 4 trials in the error clamp phase, and Asymptotic Error Clamp, defined as the final 4 trials in the EC phase, after participants reach asymptotic performance. In each phase, the average adaptation index measured across all EC trials (3-4 trials) was taken as the outcome measure for each participant. Blocks within each task are reported in Fig. 1 for reference.

Because adaptation is an error-based learning process, we performed a mixed model analysis of the same form using trajectory errors in the dynamic adaptation task as outcome measure to test for potential confounding effects of differences in errors experienced during task execution. As above, the between subject factors included robotic device and session. The experimental phases for this analysis included: Baseline, defined as the last four trials of the motor performance task, Early Counter-Clockwise and Late Counter-Clockwise, defined as the first and last four trials in block 2 and 7 of the dynamic adaptation task, respectively, After Effects defined as the first four trials of block 8, Early Relearning and Late Relearning defined as the first and last four trials of the re-introduced counter-clockwise curl force condition, respectively, and Early Clockwise, defined as the first four trials of the clockwise curl force condition.

In both cases, experimental phases were defined to capture key features of expected behavior during force field transitions and asymptotic performance. For adaptation index data, increases in the Early Counter Clockwise, Late Counter Clockwise, After Effects and Relearning phases compared to Baseline would all be indicative of adaptation. We expect the force-field reversal to cause an initial washout of adaptation to the counter-clockwise force field, that rebounds after a few trials, consistent with spontaneous recovery. As such, we expect to see no significant difference from baseline in the Initial Spontaneous Recovery phase, and significantly greater adaptation from baseline in the Asymptotic Spontaneous Recovery phase. For trajectory error data, decreases between the Late counter clockwise and Early counter clockwise phases (i.e. error reduction) would demonstrate a learned response to the dynamic perturbation, while decreases in the After Effects phase from the Baseline phase would signify after effects, indicative of adaptation.

We additionally tested for differences in behavior measured between devices in the Random Force task. We used the same mixed model structure described above, with between-subject factors robotic device and session, and within-subject factor experimental phase. We defined 8 experimental phases of interest corresponding to average behavior measured in blocks 2-9 of the random force task, for both trajectory errors and adaptation index measured across the whole task. Because the perturbations alternate directions, we used the absolute value of our trajectory error metric for this analysis. We expect to see significant increases in trajectory errors during force application, but no significant changes in adaptation index in any phase, as no significant adaptation should occur.

For all mixed-models we used a full factorial design that included the main effects of Experimental phase, Session, and Robotic Device, as well as all interaction terms between fixed factors. When the mixed model returned a significant effect of any fixed factor, post-hoc Tukey test were used to quantify the effect on behavior.

##### b) Mixed model analysis of EMG

To establish expected patterns of co-contraction during each task (dynamic adaptation, random force), we performed a one-way repeated measures ANOVA on antagonist muscle activation (ECU and FCR) and on global activation. For both tasks, we expect to see an initial increase in co-contraction in response to the unexpected perturbations. In the dynamic adaptation task, co-contraction should decrease to baseline levels as individuals adapt to produce the ‘correct’ activation pattern to reject perturbations and reduce wasted activation and metabolic costs [19], [20]. In the random force task, we expect co-contraction to decay gradually and remain elevated compared to baseline at the end of the task, as participants continue to co-contract to reduce perturbations, indicative of impedance control [12].

Experimental phases of interest were defined as follows; Baseline 1, defined as the last block of the motor performance task, Early Dynamic Adaptation and Late Dynamic Adaptation, defined as block 2 and 7 of the dynamic adaptation task, respectively, Baseline 2, the second zero force block of the random force task, and Early Random Force and Late Random Force, defined as block 3 and 8 of the random force task, respectively. For each participant, the average activation for all valid trials within each experimental phase was taken. When the ANOVA returned a significant effect, post-hoc Tukey test were used to quantify the effect of experimental phase on EMG activation. We additionally performed a control analysis of absolute trial velocity (max and mean) measured in these same experimental phases, to confirm that any effects observed in our main analysis of EMG activation were not attributable to changes in movement kinematics.

##### c) Association between excess error reduction and co-contraction

To determine the relationship between co-contraction and excess error reduction (i.e. the portion of trajectory errors not explained by adaptation) we performed a correlation between participant’s residual error reduction and their EMG measures of co-contraction at steady state compared to baseline. We additionally performed a linear regression between group average excess error reduction and average EMG measures of co-contraction across all tasks. For the group level analysis, data were binned in blocks of four trials, corresponding to two flexion and two extension trials, to reduce the influence of noise. For both group level and subject level analyses, we investigated the relationship between excess error reduction and co-contraction defined in pure antagonist muscles (FCR and ECU), as well as our global activation measure. We additionally investigated the relationship between co-contraction and alternate behavioral metrics (trajectory errors and adaptation index) to determine the specificity of our results.

## III. Results

### A. Mixed model analysis of behavior

Adaptation index and trajectory error data measured across all tasks are shown in Fig. 3 with the grey highlighted at the bottom of the figure indicating experimental phases considered for statistical analysis.

**Fig. 3.**
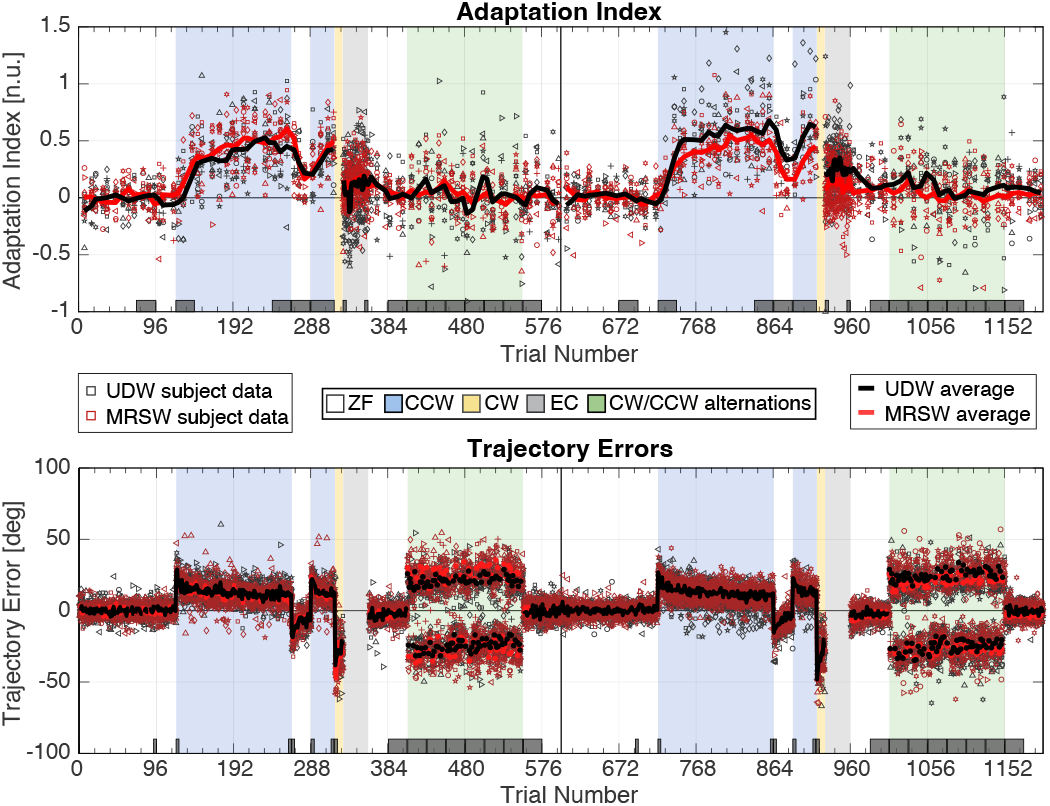
Behavioral Results. Top: Individual subject level data are reported for each device and each session (Session one: trials 1-600; session two: trials 601-1200) via marker outlines. Participants were represented with the same marker between sessions and across behavioral metrics. For visualization only, we applied a moving average of 2 trials to group average adaptation index data to smooth variability associated with intermittent sampling. Grey bins at the bottom of each figure highlight the experimental phases used in the mixed model analyses.

#### 1) Adaptation Index

The mixed model fit to adaptation index measured in the dynamic adaptation task 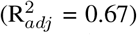 returned a significant main effect of experimental condition (*p* < 0.001) but no significant effect of Robot (*p* = 0.68), Session (*p* = 0.77), or any interaction terms. Model parameter estimates showed that adaptation index was significantly greater in all experimental phases compared to baseline (*p* < 0.001) except for the initial error clamp phase, in line with effects predicted by adaptation and spontaneous recovery.

The model fit to adaptation index in the random force task had a poor fit 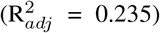, as expected given that no adaptation should occur in this task. The model returned a significant main effect of Experimental condition (*p* = 0.011) and Session (*p* = 0.0435), but no significant effect of robot (*p* = 0.79), nor any interaction terms. Post-hoc Tukey testing returned no significant difference between the baseline block and any experimental phase, in line with no significant adaptation with task execution. The main effect of session was driven by greater adaptation index in session 2 compared with session 1 (Session 1 = 0.0033 ±0.033 deg; Session 2 mean = 0.0914 ±0.033 deg), which suggests that washout of adaptation in session 2 was slower compared with session 1.

#### 2) Trajectory Errors

The mixed model fit to trajectory errors in the dynamic adaptation task 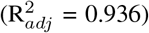 returned a significant main effect of experimental condition (p <0.001), and no significant effect of Robot (*p* = 0.34), Session (*p* =0.63), or any interaction terms. Post-hoc Tukey testing returned significant decreases in error between the early and late counter clockwise phases, indicative of learning, and significant after effects consistent with adaptation.

The mixed model fit to absolute trajectory errors in the random force task 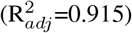 returned a significant main effect of experimental condition (*p* < 0.001), and no significant effect of Robot (*p* =0.928), Session (*p* =0.782), nor any interaction terms. Trajectory errors in all six blocks in the random force condition were significantly greater than baseline, but there were no significant after effects, nor significant change in error across the task.

#### 3) Corollary Analyses

Because our force field is velocity dependent, we performed a mixed model analysis on mean velocity measured in the same experimental phases for adaptation index and trajectory errors used above. This analysis returned no significant effect of robot nor interaction term that could confound comparisons between devices. To test for effects of device on model estimation, we used a mixed model analysis to compare the distributions of parameter estimates for a two-state model fit to subject-level behavior, that included factors Device and Session. Our results showed no significant differences in the distribution of parameter estimates for the two state model fit to either device. As such, two-state model estimation used for excess error estimations were not considered significantly different between devices.

### B. Mixed model analysis of EMG

Given that device had no significant effect on behavior, we collapsed EMG measured across devices and performed three one-way ANOVAs to test for the effect of experimental phase on EMG measures of co-contraction taken in antagonist muscles (ECU, FCR) and across all muscles (global activation).

The model fit to the ECU data 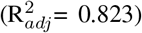 identified a significant main effect of experimental phase (*p* < 0.001). In line with our expectations, post-hoc Tukey testing identified significant increases in activation compared to baseline in the early dynamic adaptation phase, and early and late random force phases (mean ±standard error; Baseline 1: 82.39 ±18.78; Early DA: 186.48±18.78; Early RD: 180.27 ±18.78; Late RD: 144.56 ±18.78). There was no significant difference between early dynamic adaptation and early random force conditions, nor between baseline 1, Late dynamic adaptation, and baseline 2 (Late DA: 102.56±18.78; Baseline 2: 111.90 ±18.78). A contrasts on parameter estimates indicated that activation across the dynamic adaptation task decreased more than activation across the random force task (*p* =0.011). The model fit to the FCR data 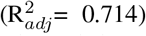 and global activation data 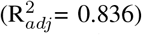 both showed the same patterns of significant effects as the ECU.

Repeated measures ANOVA of the max and mean trial absolute velocity measured these same experimental bins showed that there were no significant change in velocity across each task, nor between dynamic conditions in any experimental phase. As such, change in velocity can not account for the observed differences in activation reported above.

In sum, our measures of EMG activation show expected patterns of co-contraction in the dynamic adaptation and random force task, consistent with those reported in the literature for similar task paradigms [12], [19], [20].

### C. Association between excess error reduction and co-contraction

Fig. 5 shows the group level association between EMG measures of co-contraction and excess error reduction, defined by the squared residual error between measured trajectory errors and predicted errors from a two-state model of adaptation. The two-state model fit to average adaptation index data returned model parameters *A_f_* = 0.91689, *A_s_* = 0.99306, *B_f_* = 0.02878, *B_s_* = 0.008143, and fit trajectory errors across all tasks with an 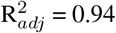. Across all tasks, a significant portion of the variability in excess error reduction was described by EMG measures of co-contraction. The linear model fit between average antagonist muscle activation and excess error reduction had an 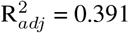, p < 0.001 across both tasks, an 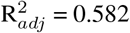, p < 0.001 in the dynamic adaptation task, and an 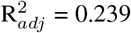, p < 0.001 in the random force task. Comparison between excess error reduction and average global activation, as well as independent antagonist muscle activations (ECU and FCR) yielded similarly significant (all p < 0.001) associations across all tasks.

**Fig. 4.**
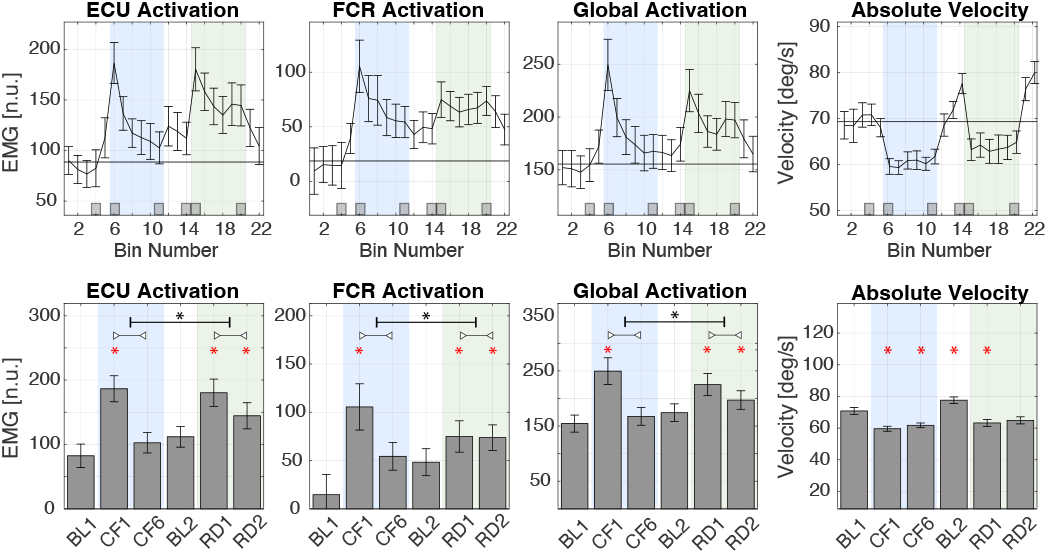
EMG Results. Top: Average EMG activation measured in each block across all tasks in session 1 (column 1-3), and the corresponding max absolute trial velocity (column 4). Grey bins denote experimental phases used for statistical analysis. Bottom: Bar plot of average behavior measured in each experimental phase included in our one-way ANOVA analysis (baseline: BL, dynamic adaptation: DA, random force: RD). Red asterisks denote changes in activation that are significantly greater than baseline 1. Post-hoc comparison of change within each task that reached significance are shown via black asterisks over a horizontal bar.

**Fig. 5.**
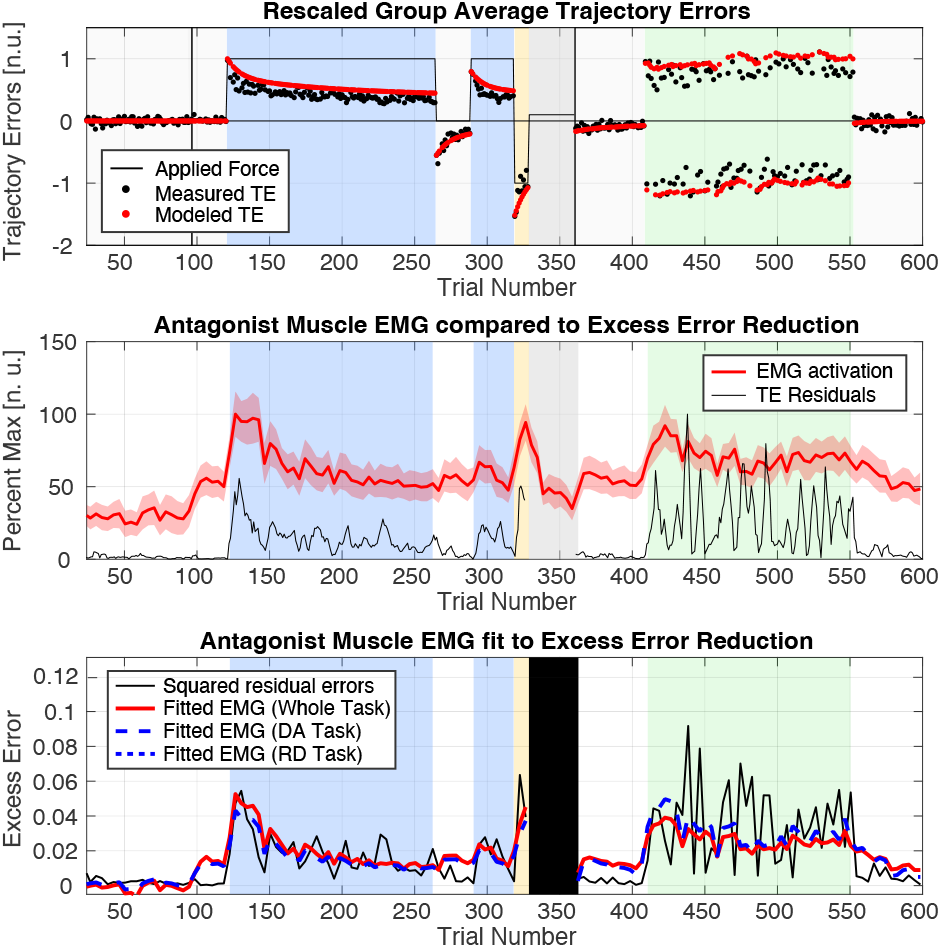
Relationship between EMG measures of co-contraction and excess error reduction measured at the group level. Top: Model predicted trajectory errors compared to rescaled group level trajectory error data. Measured errors (black) are in excess of model predictions (red). Middle: Overlay of average EMG antagonist muscle activation (red, mean and standard error) and squared residual errors (black) across all tasks. Both data sets are smoothed with a 4 bin moving average. Bottom: Average antagonist muscle EMG activation fit to squared residual trajectory errors, fit across all motor tasks (red) and separately within the dynamic adaptation (DA) and random force (RD) tasks (blue dashed lines).

Next, we investigated the association between EMG activation measured in the dynamic adaptation task to excess error reduction quantified at the end of the constant perturbation phase of the dynamic adaptation task (block 7), in which we previously observed excessive error reductions not explained by our models of adaptation (Fig. 6). The two state model fit to subject-specific data had an average fit of R^2^= 0.536 ±0.023, range [0.391, 0.723] to adaptation index data, and R^2^ = 0.625 ±0.091, range [0.299, 0.810] to trajectory errors.

**Fig. 6.**
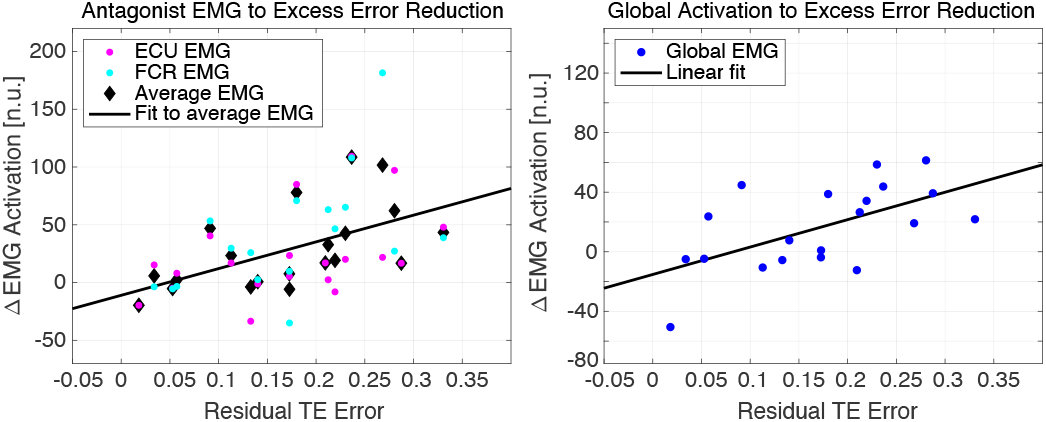
Linear regression between residual trajectory errors (excess error) from subject-specific model estimations to change in EMG activation in pure antagonist muscles (left) and global muscle activation (right). Change in EMG activation was measured with respect to baseline levels of activation measured in the zero force condition. Positive residual errors are associated with greater error reduction compared to model predicted values (residual error = modeled error - normalized error). On the left, ECU muscle activations are shown in magenta (R = 0.509), FCR in cyan (R = 0.564) and the average across muscles is shown in black (R = 0.617). On the right, global activation is shown in blue (R = 0.59). For both figures, the linear regression line is shown in black.

Excess error reduction was significantly correlated with change in EMG activation compared to baseline within each antagonist muscle (FCR: *r* = 0.564, *p* = 0.0184; ECU: *r* = 0.509, *p* = 0.022), across both muscles (*r* = 0.617, *p* = 0.0038), and to global EMG activation (R = 0.59, p = 0.006). All analyses returned a positive association between larger EMG activation and greater error reduction compared with that predicted by models of adaptation.

To test the specificity of these results, we additionally investigated the association between measured trajectory errors and modeled adaptation in this phase to changes in EMG activation. This analysis returned no significant associations within either muscle nor across muscles for trajectory errors (*p_ECU_* = 0.70, *p_FCR_* = 0.59, *p_Global_* = 0.15), change in trajectory error (*p_ECU_* = 0.9265, *p_FCR_* = 0.083, *p_Global_* = 0.75) nor magnitude of adaptation (*p_ECU_* = 0.40, *p_FCR_* = 0.49, *p_Global_* = 0.77).

Taken together, these analyses establish that muscle activation associated with co-contraction explains a significant portion of the variance in measured trajectory errors that is not explained by measures of adaptation. In both analyses, greater levels of co-contraction, approximated by EMG activation in antagonist muscle groups, were significantly associated with error reductions in excess of those predicted by models of adaptation learning.

## IV. Discussion

We performed this study with two aims: to validate behavior measured with the MR-SoftWrist, and to investigate the concurrent use of co-contraction during dynamic adaptation of the wrist. To address these aims, we used a two-session counter-balanced experimental design. For both sessions, participants performed the same task schedule, consisting of a zero force task, a dynamic adaptation task, and a random force task. Participants used either the MR-SoftWrist or the UDiffWrist for task execution in session one, and the other robotic device for session two, conducted 3-4 weeks apart. For all tasks, muscle activation of the flexor carpis radialis (FCR), flexor carpis ulnaris (FCU), extensor carpis radialis (ECR), extensor carpis ulnaris (FCU) was collected using surface EMG.

To validate measurements of motor behavior collected via the MR-SoftWrist, we compared outcomes measured with the MR-SoftWrist to those measured with the UDiffWrist, a functionally equivalent non MRI-compatible wrist robot. Mixed model analysis showed no significant main effect of Device on behavior (adaptation index, trajectory errors, velocity) in either the dynamic adaptation or the random force task. Moreover, comparison of the two-state model fit to subject-level adaptation index data showed no significant difference in model estimation between devices. Together, these results validate that adaptive behavior measured with the MR-SoftWrist is not unduly influenced by the compliance of the device.

In our previous studies, we observed a lower end-magnitude of adaptation (study 1: mean ±s.e.m.: 0.355 ±0.0257, after 144 trials; study 2: 0.43 ±0.05, 180 trials) than is commonly observed in reaching tasks (0.5-0.7) [8], [9], [15]. However, participants in this study achieved a greater end magnitude of adaptation (0.59 ±0.046). Increased adaptation index may be due to modifications in experimental design, including a new handle design, slightly stiffer error clamp trials and slightly lower forces. As force modifications were minor, we believe these changes are likely due to the change in handle design. Our previous design required subjects to grasp the handle with a closed fist, necessitating activation of extrinsic muscles in the forearm that may increase co-contraction and greater use of impedance control. However, experiments controlling for other factors are needed to confirm our speculation.

Next, we aimed to establish patterns of concurrent cocontraction during dynamic adaptation of the wrist. We quantified co-contraction via EMG activation measured in pure antagonist muscles (FCR for extension trials, ECU for flexion trials) and as global activation across all relevant wrist muscles. Analysis showed initial increases in activation in both tasks, followed by a significantly greater decrease in activation to baseline levels in the dynamic adaptation task. These results are consistent with established patterns of cocontraction associated with impedance control, in that activation remains elevated in unlearnable dynamic conditions, but significantly decreases in constant force fields as subjects adapt and the correct muscle patterns are learned for more efficient perturbation rejection. These results were consistent across all EMG definitions tested, and serve to validate that our measures capture concurrent co-contraction processes.

Finally, we investigated the contribution of co-contraction to error reductions not attributable to adaptation. We aimed to determine if variance in residual errors between measured trajectory errors and those predicted by a two-state model of adaptation was explained by our EMG measures of co-contraction. Linear regression analysis showed that the time course of average antagonist muscle activation explained a significant portion of variance in average excess error reduction measured across the whole task (39%), and within the dynamic adaptation task (58%) and the random force task (24%). Correlational analysis of behavior measured at steady state in the dynamic adaptation task showed that excess error reduction was significantly associated with greater EMG activation (i.e. greater co-contraction) across all EMG metrics (ECU, FCR, global). Importantly, this association was specific to excess error reduction, as there were no significant association between EMG activation and alternate measures of trajectory error or adaptation. These results support our hypothesis that for dynamic perturbations of the wrist, reductions in trajectory errors in excess of those explained by adaptation are largely attributable to concurrent impedance control (co-contraction).

In short, this study establishes validity of the measurements of motor behavior obtained using the MR-SoftWrist, and establishes the concurrent involvement of impedance control in the response to force perturbations applied via this device. These findings advance our ability to investigate via fMRI the neural substrates responsible for each neuromotor control process involved in dynamic adaptation for tasks involving the hand and wrist.

## Acknowledgment

We acknowledge support from the American Heart Association (award no. 17SDG33690002), and from the National Science Foundation (award no. 1943712).

